# Computational modelling of the long-term effects of brain stimulation on the local and global structural connectivity of epileptic patients

**DOI:** 10.1101/728576

**Authors:** Emmanouil Giannakakis, Frances Hutchings, Christoforos A. Papasavvas, Cheol E. Han, Bernd Weber, Chencheng Zhang, Marcus Kaiser

## Abstract

In patients with drug resistant focal epilepsy, targeted weak stimulation of the affected brain regions has been proposed as an alternative to surgery. However, the effectiveness of stimulation as a treatment presents great variation from patient to patient. In this study, brain activity is simulated for a period of one day using a network of Wilson-Cowan oscillators coupled according to diffusion imaging based structural connectivity. We use this computational model to examine the potential long-term effects of stimulation on brain connectivity. Our findings indicate that the overall simulated effect of stimulation is heavily dependent on the excitability of the stimulated regions. Additionally, stimulation seems to lead to long-term effects in the connectivity of secondary (non-stimulated) regions in epileptic patients. These effects are correlated with a worse surgery outcome in some patients, which suggests that long-term simulations could be used as a tool to determine suitability for surgery/stimulation.

## Introduction

Pharmaceutical drugs that can pass through the blood-brain-barrier lead to changes in the whole brain, which can result in severe side effects. Moreover, for many patients these traditional approaches do not work well in treating the symptoms of brain network disorders. Instead, targeted approaches that only *directly* affect a small number of brain regions have been proposed. These techniques range from localised opening of the blood-brain-barrier through focused ultrasound, to invasive and non-invasive brain stimulation, and, when no alternative options are suitable, to surgical removal of brain tissue. The problem then is to choose the right set of target regions for individual patients to maximize treatment effects and to minimize side effects.

Parkinson’s disease and epilepsy are diseases where targeted approaches are already routinely used, when drug treatment is insufficient. For focal epilepsy, where medication is ineffective, resective surgery of the affected regions is often used as a treatment. However post-operative seizure remission is around 50-70% (1, 2). The reoccurrence of seizures after surgery could be due to incomplete removal of the required target regions (3) or due to surgery causing remaining brain regions to become new starting points for seizures. For the latter option, it will be crucial to develop computer models of long-term effects of interventions.

The same challenge occurs for brain stimulation in epilepsy patients where no tissue is resected but where the stimulation of a target region, with reduction of epileptogenic activity in that region, could potentially cause other non-stimulated regions to become starting points for seizures. Targeted brain stimulation in epilepsy could include deep brain stimulation (DBS), optogenetic stimulation (4) (www.cando.ac.uk), and non-invasive techniques (transcranial current stimulation, TCS; transcranial magnetic stimulation, TMS). The effectiveness of those methods varies (5) and when it comes to TCS– one of the non-invasive methods–there are of contradictory results concerning its efficacy for treating epilepsy (6-11). Across several studies of TCS, 67% of studies show a decrease in seizures after stimulation (6). Some of these studies covered in that article examined the effects of stimulation only for a short period after the end of the stimulation session (hours) without a subsequent follow up. Thus, it is possible that the long-term efficacy of TCS is not as high as 67%.

One of the main concerns with TCS is whether the effects of stimulation would remain after the stimulation has ended (12). Some studies have shown that the positive physiological effects of stimulation can outlast the stimulation session for a long period while others have shown diminishing effects after the stimulation session has ended. Specifically (13-15) have observed positive post-stimulation effects lasting for a period of 2, and more than 4 months respectively. On the other hand (16) observed anti-seizure effects for a period of 48 hours after stimulation but also a clinically significant reduction of those effects during a subsequent period of 4 weeks. To use computational models to assess the effect of brain stimulation, it is therefore necessary to observe long-term changes.

At the moment, computational studies have only examined the short-term effects of TCS, i.e. during stimulation (17-21). Two computational studies have used neural mass models (22, 23) to examine the immediate effects of stimulation on the activity of the stimulated areas. Notably, one study used modified Wilson-Cowan model to study effects a few minutes after anodal or cathodal stimulation (23). The aforementioned studies did not account for plasticity in their models, and so did not investigate the effects of stimulation on brain connectivity. The only computational study to our knowledge that does examine the effects of neurostimulation on brain connectivity (24) focuses on DBS instead of TCS and examines Parkinson’s disease instead of epilepsy with the aim of identifying optimal stimulation locations.

In this study, we will observe long-term changes after initial stimulation in terms of both structural connectivity changes and changes in local and global network dynamics. We focus on connectivity changes as only such changes at the structural level can explain the behaviour of networks a long time after the initial stimulation and thus could explain the final outcomes of treatment (25). We find that simulated effects of brain stimulation differ between epilepsy patients and healthy subjects, (2) stimulation leads to distinct long-term connectivity changes in non-stimulated regions, and that (3) these indirect effects after stimulation are more informative for outcome predictions (using surgery outcome as a basis for prediction) than direct effects that are observed during the stimulation.

## Results

For the purposes of this study, we can group our simulation subjects in three categories, according to the global connectivity data and model used:

1. Healthy subjects: The global connectivity data were derived from healthy individuals and the simulation was performed using a model where no epileptogenic (particularly excitable) regions are present.
2. Epileptic subjects: The global connectivity data were derived from individuals suffering from left temporal lobe epilepsy and the simulation was performed using a model where the stimulated regions were modelled as epileptogenic (highly excitable)
3. Control subjects: The global connectivity data were derived from individuals suffering from left temporal lobe epilepsy but the simulation was performed using the “healthy” model, where the stimulated regions are not distinct in terms of excitability from the other regions.

Our results are organized in two sections. Firstly, we simulate the effect of stimulation on the overall connectivity of the brain for each group of subjects. Secondly, we simulate the changes stimulation seems to induce in each brain region with emphasis at the stimulated regions which are most often associated with seizure generation (amygdala, hippocampus and parahippocampal gyrus).

Statistical results will be presented for the rest of the paper as: X ± Y, where X is the mean and Y is the standard deviation of the referenced dataset. All the p-values were calculated using a two-tailed t-test.

### Patients show a larger global connectivity change at the end of the stimulation

The effect of stimulation on the connections between nodes in our model follows a similar pattern in all subjects. Specifically, during the period of stimulation, the global effect measure *D*(*t*) increases steadily (Figure 1), reaching a local maximum at the end of stimulation (*t* = 2000 *s*). A first difference between the three groups can be observed at this point since the value of *D*(*t*) at the end of stimulation is on average significantly (p-value < 0.0001) greater for the epileptic subjects (2.9730 % ± 0.7301) than the healthy subjects (1.9671% ± 0.3261) .and the control subjects (1.7609% ± 0.5290). The similarity of the healthy and control groups in contrast to the epileptic group suggests that the increased excitability of the stimulated regions and not the global connectivity is the main driver of the changes of the global effect measure. Indeed the global connectivity on its own seems to make the healthy subjects more excitable, since the values of *D*(*t*) were slightly higher for the healthy than the control group (although the difference was not statistically significant).

**Figure 1.**
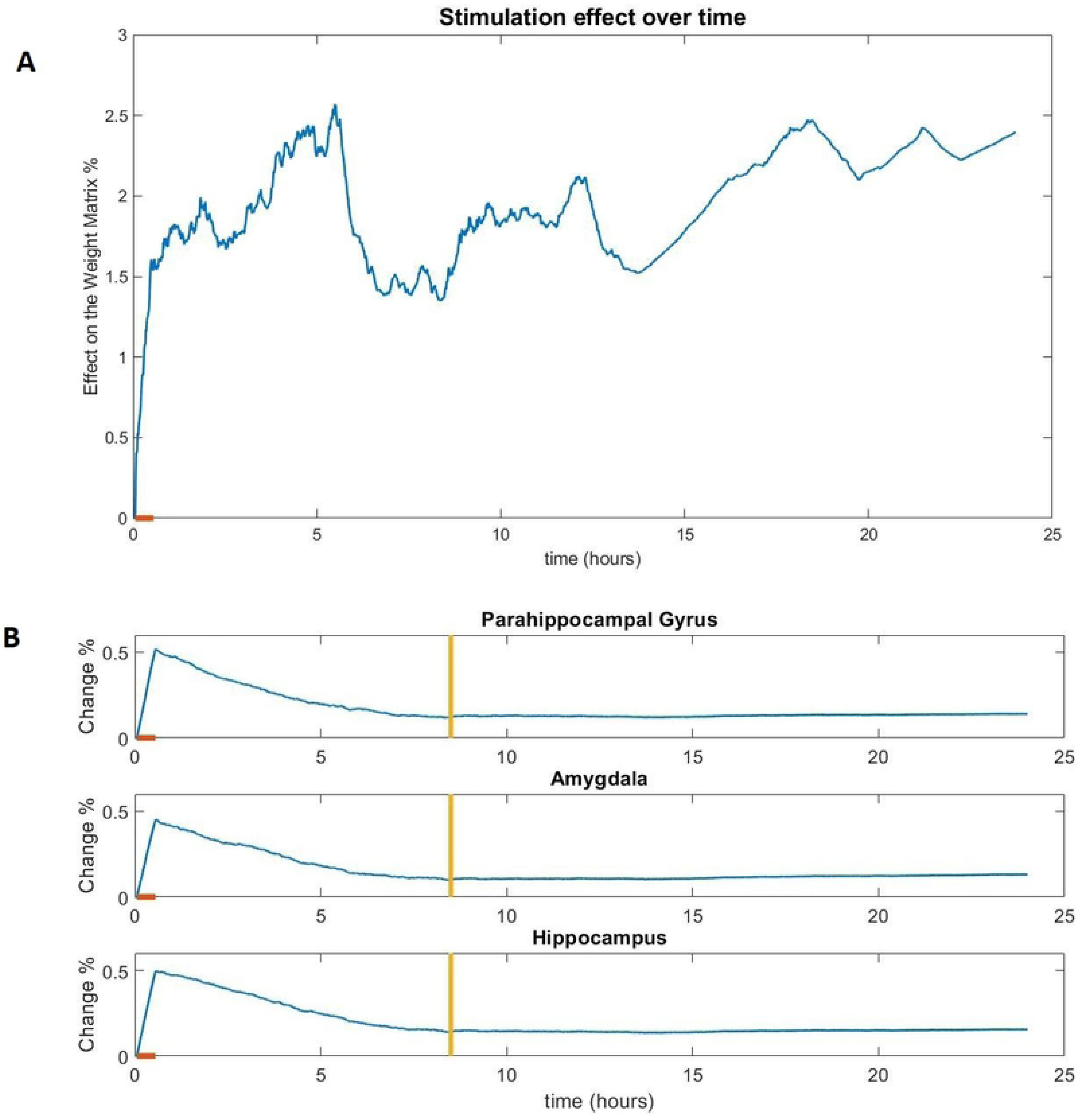
The effect of stimulation (difference from the non-stimulated version) on the global connectivity (A) and the connectivity of the stimulated regions (B) of a healthy subject. The orange line on the x-axis notes the duration of the stimulation session.

After the end of stimulation, the global effect *D*(*t*) keeps fluctuating for the remainder of the simulation with a clear increasing trend in the majority of subjects. The rate of this increase varies greatly from subjects to subject and it was calculated as the rate *r* = *D*(*t*_0_) /*D*(*t*_1_), where *t*_0_ = 2000*s* is the end of the stimulation session and *t*_1_ = 24*h* the end of the simulation. For all subjects the value *d* varies greatly (0.5846% ± 0.2751) and we can also observe a small difference (statistically insignificant) between the values of healthy subjects (0.5358% ± 0.2128), the similar values of control subjects (0.5372% ± 0.1609) and the slightly greater values of epileptic subjects (0.6328 % ± 0.2533) which is not statistically significant. Thus, the differences between the groups are attributable to different effect of stimulation and not the post-stimulation change in connectivity.

Finally, the correlation between the development of the global effect measure of the control subjects and the equivalent epileptic subjects (using the same global connectivity data), is significantly (p-value = 0.0476) higher (0.7747 ± 0.1102) than the correlation between random pairs of control and epileptic subjects (0.6199 ± 0.3213). This, suggests that although the scale of change is mainly determined by the excitability of the stimulated regions, the exact global connectivity does (at least partially) determine the development of the global effect measure.

### Patients show a larger change in local connectivity of stimulated regions during but not after stimulation

In the regions that received direct stimulation (amygdala, hippocampus and parahippocampal gyrus), the effect on the connectivity was most prominent during the period of stimulation, resulting in a constant increase of the local effect measure d_k_(*t*) in all three regions. Thus, the local measure invariably reaches a global maximum at the end of the stimulation session (*t* = 2000 *s*). As with the global connectivity, the effect on the epileptic subjects is greater than the effect on the other two groups (p-value < 0.0001 for all three regions). Specifically, the average effect for all three regions on a healthy subject is 0.4746 % ± 0.0509, in a control subject is 0.3853 % ± 0.0427 and on an epileptic subject is 1.0794% ± 0.0264.

A difference from global connectivity is that in this case the difference between healthy and control subjects is clearly significant (p-value < 0.0001). This suggests that the brain connectivity of epileptic patients conditions the epileptogenic regions to be less excitable than in healthy individuals. Of course, the internal connectivity that makes these regions highly excitable mask this effect as we observed from the metrics of the epileptic group. Still, this finding seems to suggest that the inter-regional connectivity of epileptic patients tends to limit the excitability of epileptogenic regions.

After the end of the stimulation session, the local measure *d*_*k*_(*t*) changes similarly in the healthy/control groups but very differently in the epileptic group.

In the healthy/control subjects, the end of the stimulation session is followed by a slow decrease in the value of the local effect *d*_*k*_(*t*). Around 8 hours after the end of the stimulation session, the difference measure stabilizes at *d*_*k*_(*t*) ≈ 0.1 %, for all three regions (Figure 1), for a representative subject. The local effect measure *d*_*k*_(*t*) of a region is considered to be stabilized at time *t* if the Coefficient of variation of the values of *d*_*k*_(*t*) for the 5 minutes prior to *t* is less than 0.3. After that point, there may be some small oscillation in the value of *d*_*k*_(*t*) but the change is minimal.

There is much greater variation in the epileptic subjects, both between the regions of the same subject as well as between equivalent regions of different subjects (Figure 2). Immediately after the end of the stimulation session and for a period lasting 5-6 hours, the local effect *d*_*k*_(*t*) is sharply (more than in the healthy/control subjects) decreasing for all 3 regions. With the exception of two subjects where there is a short increasing period in the values of the amygdala and the hippocampus, *d*_*k*_(*t*) is strictly decreasing during this period for all three regions of every subject. It should be noted that in almost all the epileptic subjects (91%), the connectivity of the parahippocampal gyrus is behaving differently than the connectivity of the other two regions. The local effect (measured by *d*_*k*_(*t*)) on the parahippocampal gyrus is diminishing faster than the equivalent measures of the other two regions, reaching values close to zero at the end of this first period.

**Figure 2.**
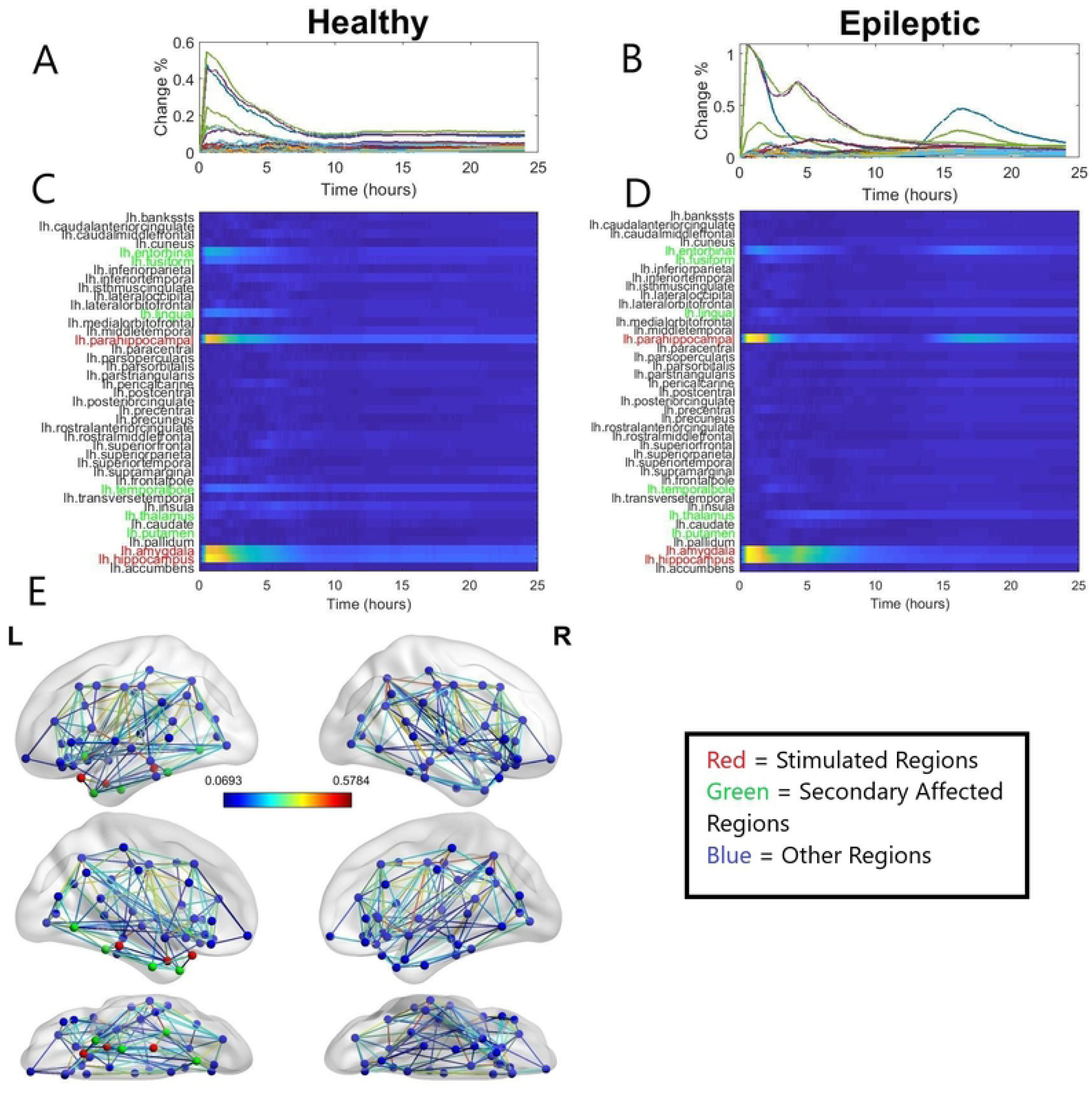
A - D: The local effect on the connectivity of all the left hemisphere regions of a healthy (A, C) and an epileptic subject (B, D). The greater effect of stimulation on the epileptic subject can be observed as well as the influence on secondary (non-stimulated) areas in both subjects. Also, the difference between the effects on the Parahippocampal Gyrus and the other regions can also be seen on the epileptic subject. L-R: The brain network of the epileptic patient. The red nodes indicate the stimulated regions and the green nodes indicate the secondary affected regions. The colour of the edges indicates the strength of the connection.

For the remainder of the stimulation, each subject presents different behaviour and the various stimulated regions also present differences in each subject. In 50 % of the subjects the local effect on the parahippocampal gyrus remains at the low levels it reached in the end of the decrease period (1 – 5/6 hours) with some minimal increases. In the remaining 50 % the local effect on the parahippocampal gyrus starts increasing at some point between 8-12 hours after the end of stimulation and continues to increase for the remainder of the simulation reaching values comparable with those of the other two regions. The other two regions (amygdala and hippocampus) behave almost identically in each subject. After the end of the first period of decrease the local effect measures of these areas stabilize in 50 % of the subjects and decrease very slowly in 33.5% of the subjects for the remainder of the stimulation. In the remaining 16.5 % of the subjects, the local effect measure increases for a period of 1.5 -2 hours until it reaches values much higher than those of the other subjects (*d*_*k*_(*t*) ≈ 0.55), after that point the effect on those areas begins to slowly decrease.

At the end of the simulation, we can observe that the final values of d_k_(t) for the epileptic subjects (0.1412% ± 0.0882) are slightly greater than those of the healthy subjects (0.1165% ± 0.0275) which in turn are slightly greater than those of the control subjects (0.1037% ± 0.0400) in the regions that received stimulation. Still that differences are not statistically significant. This implies that the initial difference between healthy/control and epileptic subject does not lead to a long-term difference in the stimulation effects.

### Some non-stimulated regions show local connectivity changes after the end of the stimulation

The effects of stimulation can be seen not only on the internal connectivity of the regions that are stimulated directly but also on the connectivity of other brain regions that receive no direct brain stimulation (Figure 2).

Specifically, in all groups, the local effect *d*_*k*_(*t*) of several regions starts increasing and reaches a peak shortly after the end of the stimulation session. It should be noted that the change in those regions does not absolutely coincide with the stimulation session, rather it happens shortly afterwards, possibly due to the time delays. Moreover, unlike the stimulated regions where a difference can be observed between healthy/control and epileptic subjects, no such difference can be observed in the values of those secondary regions.

After this initial increase, the local effect on all secondary regions usually decreases and seems to stabilize after a period of about 8 hours. For the majority of subjects (75%) the values that the difference measures have at this point will be very close to the values they will have at the end of the stimulation. In most cases, the final value of the effect measures for those regions are very close to the values of the other non-stimulated regions that were not affected by the stimulation, but in some cases the final values for some of those secondary regions (especially the entorhinal cortex) are much closer to – and in some cases higher than - the values of the stimulated regions. Interestingly, in some epileptic subjects (25 %) the local effect measure of some secondary regions began to suddenly increase hours after the stimulation session when they were apparently stabilised for some time. This may be evidence for long-term effects that cannot be predicted from the initial response to stimulation.

It should be noted that as with the stimulated regions, all of the secondary regions refer to the left hemisphere of the brain.

Several factors could explain why those regions in particular were affected. Specifically, those regions were characterized by increased connectivity with the stimulated regions as well as small Euclidian distance from the stimulated regions. Additionally, the effect the connections with the stimulated regions seemed to be greater than average (increased connection weights). Finally, the Jaccard index (common neighbours) of the affected regions and the stimulated regions was higher than in regions that were not affected. Moreover, the frequency of excitation among the six regions that were excited is correlated with the aforementioned metrics. For example, the entorhinal cortex that was affected in 88% of the subjects scores higher in all the metrics (connectivity, Jaccard index, etc) than the putamen which was excited in less than 10% of the subjects. A ranking of all the regions according to those metrics as well as the corresponding absolute values are presented in the supplementary information (Table S1).

### Long-term changes, long after stimulation, are more informative of treatment outcomes

The epileptic patients from our dataset had received respective surgery of the seizure causing brain regions and the outcome of these surgeries was known for a number of them (17 subjects). The surgery carried out involved an amygdalohippocampectomy, resecting areas from the three regions that we have stimulated in our model. The observed outcome in terms of being seizure-free after surgery might of course be different from an outcome after stimulation. Nonetheless, we wondered whether our framework, might show some link with the outcomes after surgery, potentially providing us with a tool for predicting surgical success. In particular, we explored which timeframe within our simulation would be most informative in terms of predicting outcomes.

We found that an increased effect in the secondary regions that was observed in the epileptic group was correlated with a worse outcome of resective surgery: Epileptic subjects who presented a long-lasting effect on secondary regions after stimulation within our model, i.e. higher values of the local effect measures compared with other non-stimulated regions *at the end of the simulation*, were on average less likely (3.225 ± 1.220 on the ILAE classification scale) than those who did not present such effects (2.011 ± 1.110) to benefit from surgery (p-value = 0.0484, Cohen’s d = 1.042).

We next tested the outcome predictions depending on the time within the simulation (Figure 3). For this, we observe the local effect of stimulation on directly affected areas (the three targets) and indirectly affected areas. First, effects for secondary regions are more informative in terms of outcome than for the primary targeted regions. This holds throughout the observed simulation time from 1 to 10 hours. Second,

**Figure 3.**
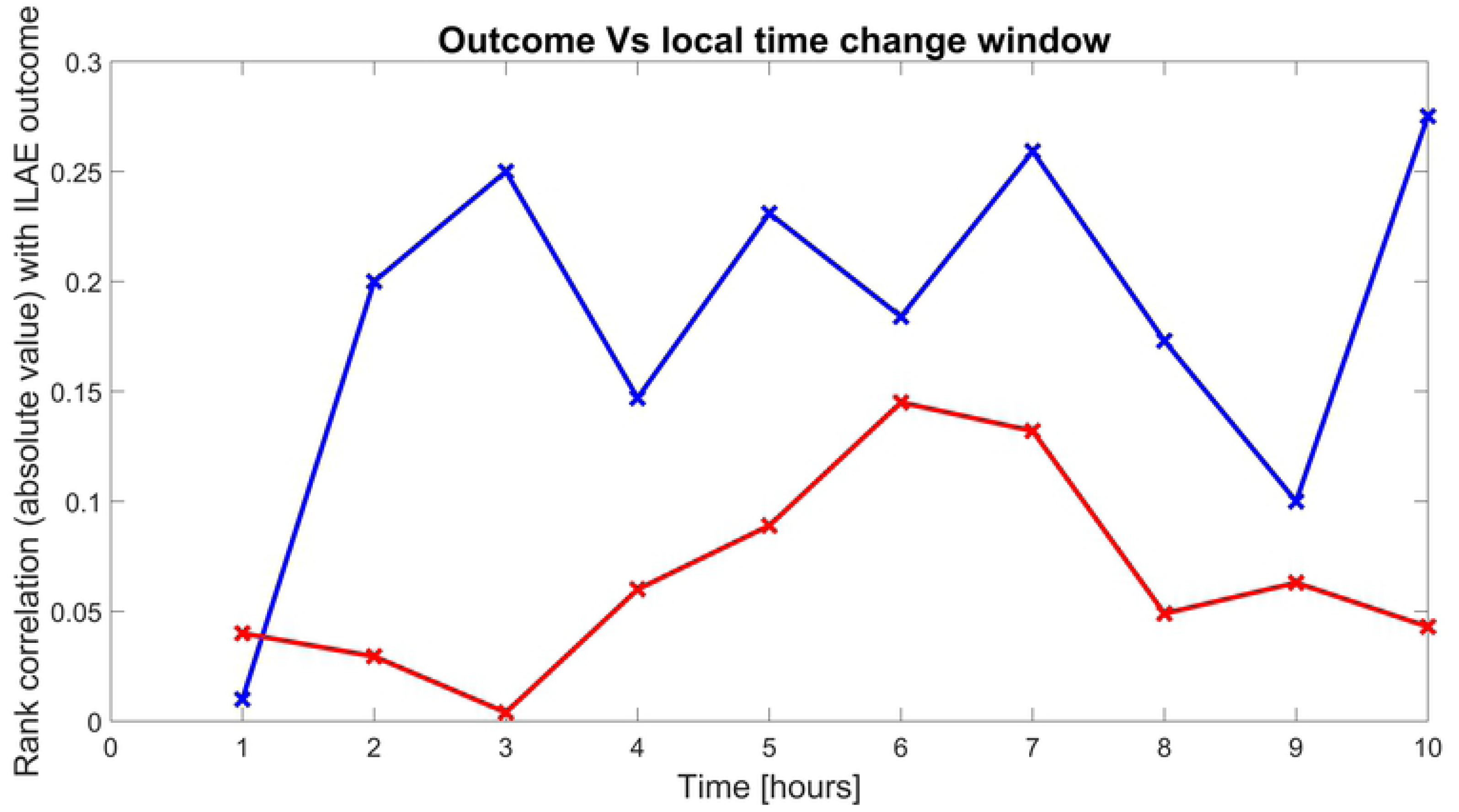
Correlation between surgery outcome and the local effect of simulation at the stimulated (red) regions and non-stimulated (blue) regions.

The effect of stimulation on the secondary regions seems to have a significantly higher correlation with the outcome of the surgery than the effect on the stimulated regions as can be seen in figure 3. Moreover, in figure 3 we can see that the effect in secondary regions is meaningful if observed hours after the initial simulation session. A greater role of secondary regions in seizure onset in the patients that show increased secondary excitation could potentially explain this correlation. Second, later time points, more than two hours away from stimulation for secondary non-stimulated regions and more than six hours away for primary stimulated regions were more informative than earlier timepoints. This could highlight that measurements several hours after the stimulation might be more useful in clinical settings to assess the likely benefit of an intervention.

The increased effect of stimulation on secondary regions could be used as a standard to determine how likely a patient is to benefit from implanted electrodes or surgery. Specifically, if our standard was to be applied as a biomarker test of suitability for surgery, it would be characterized by accuracy = 0.7059, specificity = 0.7778 and sensitivity = 0.6250, if we considered as good any surgery outcome with ILAE scores 1 and 2 and as bad any surgery outcome with an ILAE score of 3 or above (see suppl. Figure. S2).

## Discussion

We investigated the effects of simulated cathodal TCS on the brain connectivity of healthy and epileptic subjects using a network of coupled Wilson-Cowan oscillators. Our results show that stimulation affects the simulated brain connectivity—a finding that has been confirmed by experimental studies (26) —as well as a significant difference between the effect stimulation has on different groups of subjects. Moreover, the differences in the effects observed suggest that the brain anatomy of each patient may affect the long-term outcome of a stimulation session. Finally, we have observed that the effects of stimulation are not limited to the stimulated brain areas. In some patients the internal connectivity of a number of non-stimulated areas is affected by the stimulation of neighbouring areas and this seems to be correlated with a worse surgery outcome, a fact that may have some clinical significance.

Our main observation is the different behaviour of our model under the different initialisations (healthy, epileptic and control groups). In all the cases we examined, the effect of stimulation both on the internal connectivity of the stimulated regions as well as on the overall connectivity of the brain was greater on the epileptic than the healthy and control subjects which behaved similarly. This difference, combined with the observation that the effect on the non-stimulated regions (secondary regions) was similar in all groups of subjects, suggests that the increased excitability of the epileptogenic regions is responsible for the greater short-term effect of stimulation on the epileptic subjects.

Moreover, the significantly higher local effect of stimulation that was initially observed in the healthy subjects compared with the control subjects, suggests that there are indeed differences in the global connectivity of healthy and epileptic individuals and additionally indicats that the global connectivity of epileptic subjects tends to counter the epileptogenic effects of local connectivity. Finally, the long-term effects of stimulation on the internal connectivity were similar in all groups despite the initial differences, suggesting that the stimulation effect diminishes with different rates in each group.

Another finding is the great variation in the observed responses to stimulation among subjects of the same group. The extent to which the inter-regional connections change, the long-term preservation of the changes on the internal connections and the excitation of secondary regions, differed a lot from subject to subject despite the fact that the initial connectivity matrix was the only factor differentiating the model used for each subject. This fact suggests that the great variability in the effectiveness of stimulation may ultimately be caused by the differences in brain anatomy among patients. Especially given that the internal connectivity within brain areas will also differ among subjects, a fact excluded from our model as information on this was unavailable, it seems likely that the individual connectivity will be a decisive factor in determining the long-term effects of stimulation.

Moreover, the effects on the secondary regions that seem to appear without any prior indication, long after the end of the stimulation session, may indicate that effects of stimulation could appear long after the end of a session in brain regions where no stimulation was applied. In our study, we observed this phenomenon in almost 25% of the epilepsy subjects within a period of 24 hours. Still, given the lack of prior indicators for this behaviour it is possible that these sudden changes in the local effect measures could appear in more subjects or in more regions if the simulations run for a longer period of time. We examined the possibility that these sudden changes in connectivity are due to computational errors in the simulation, but the fact that the regions that present this sudden secondary excitation are almost always regions that were affected immediately after stimulation (Table 1) as well as the clinical significance of long term secondary excitation, suggest that this phenomenon is more likely attributable to the dynamics of the system and the underlying biological reality rather that to computational errors. Moreover, this phenomenon may be able to explain some of the unexpected long-term effects of TCS that appear in parts of the brain that were not stimulated. An example of this phenomenon is presented in (27), where seizures reoccur starting from a different brain region a month after an initially successful application of TCS.

**Table 1:** The non-stimulated regions that were most commonly affected in all subjects. The table shows the frequency of those effects, the local effect measure at the end of the simulation, the number of regions they are connected with and how many of these neighbouring regions are stimulated, the average effect of these stimulated regions (this was represented as the sum of the weights of the connections with stimulated regions divided with the sum of all weights) and their average Euclidian distance from the stimulated regions

| Region name | Frequency of excitation (% of subjects) | Mean and standard deviation of the final value of $d_k(t)$ | Connections with other regions / connections with stimulated regions | Average effect of stimulated regions | Average distance from the stimulated regions |
| --- | --- | --- | --- | --- | --- |
| Entorhinal cortex | 88 | 0.0766 ± 0.0360 | 6/3 | 0.5233 ± 0.0866 | 20.399 |
| Fusiform gyrus | 72 | 0.0470 ± 0.0223 | 9/2 | 0.2240 ± 0.0412 | 26.185 |
| Lingual gyrus | 44 | 0.0354 ± 0.0223 | 9/1 | 0.1283 ± 0.0302 | 49.521 |
| Temporal pole | 12 | 0.0425 ± 0.0244 | 8/1 | 0.1128 ± 0.0761 | 35.461 |
| Thalamus | <10 | 0.0714 ± 0.0529 | 12/2 | 0.2081 ± 0.0479 | 26.4036 |
| Putamen | < 10 | 0.0286 ± 0.0141 | 13/1 | 0.0357 ± 0.0101 | 26.1245 |

Finally, concerning the clinical significance of our findings, we have established a correlation between long-lasting effects of stimulation on the internal connectivity of some secondary regions and a worse surgery outcome. Specifically, we have shown that observing the long-term effect, lasting at least for several hours, of stimulation on secondary regions is more informative concerning the potential surgery outcome than observing the effects on the stimulated regions. The reason for this could be that in patients with more excitable secondary regions, these regions might still cause seizures after the primary epileptogenic regions have been removed. This correlation is not as effective as a method of predicting surgery outcome as other similar techniques (15, 28, 29) but it could be used as a secondary test to determine suitability for surgery and/or implanted electrodes. Moreover, the fact that this correlation was observed by only taking into account the intra-regional connectivity of the patients and given that the individual anatomy of each region almost certainly plays a role, it is possible that more detailed individualized simulations of this kind could predict the potential effects of surgery/stimulation in epileptic patients.

## Limitations

Our study is far from conclusive for two main reasons. Firstly, the models we used are very rough approximations of the underlying biological reality and thus, the clinical significance of our findings is far from certain. Special attention should be paid on the use of an unconventional learning rule as well as the fact that many of our constants were chosen to facilitate the simulation and thus, they may not represent the reality of biological systems. Also, local connectivity was initialised based on a previous model whereas measurements of fMRI allow for model parameters derived from subject-specific activity across brain regions (30).

Secondly, due to time limitations only one stimulation session was modelled with a subsequent resting period of 24 hours. Although our results do capture an abnormal behaviour (changes in secondary regions), it is clear that given that in many of the studies discussed in the introduction the follow up period was ranging from several days to a little less than a year, our results may not represent the behaviour of biological systems for such long periods of time.

In addition to those two main issues, it should be noted that our dataset was quite small (19 patients and 20 controls) and thus the clinical significance of our findings needs to be verified through larger datasets and experimental stimulation data. In particular, patient cohorts with brain stimulation data and simulation experiments of longer duration will be crucial to validate the predictive power of this model, since our current observations are based on surgery outcome which may differe from stimulation outcome.

## Conclusion

This study uses computational methods to examine the long-term effects of TCS on the connectivity of the brain. Our findings indicate that even small differences in the internal connectivity—and thus the excitability—of the stimulated regions can radically change the way stimulation affects the brain. Moreover, the initial connectivity between brain regions also greatly affected the way each subject behaved post-stimulation. In addition, the effect stimulation has on non-stimulated brain regions seems to be a potential biomarker of long-term treatment outcome. Finally, sudden and seemingly unprovoked changes in the connectivity hours after the effects of stimulation could explain the unexpected effects of TCS that have been observed in the past.

## Methods

### Patient data

We examined 39 subjects, 19 of whom are suffering from left temporal lobe epilepsy. The subjects were selected from the dataset presented in (31, 32). Written informed consent was obtained, signed by all participants, and conformed to local ethics requirements. The ethical review board of the medical faculty of Bonn gave IRB approval (032/08) and all experiments were performed in accordance with relevant guidelines and regulations. T1 weighted MRI scans and diffusion tensor imaging (DTI) data were obtained using a 3 Tesla scanner, a Siemens MAGNETOM TrioTim syngo (Erlangen, Germany). The T1 images were obtained using 1mm isovoxel, TR = 2500ms and TE = 3.5ms. The DTI data used 2mm isovoxel, TR = 10,000ms, TE = 91ms and 64 diffusion directions, b-factor 1000s mm-2 and 12 b0 images. In both cases FoV was 256mm.

To create the structural connectomes, FreeSurfer was used to obtain surface meshes of grey and white matter boundaries from the MRI data and to parcellate the brain into regions of interest (ROI) based on the Desikan atlas (33, 34). This process identified 82 ROIs which spanned cortical and subcortical regions (Nucleus accumbens, Amygdala, Caudate, Hippocampus, Pallidum, Putamen and Thalamus). Streamline tractography was obtained from DTI images using the Fiber Assignment by Continuous Tracking (FACT) algorithm (35) through the Diffusion toolkit along with TrackVis (36). First, we performed eddy-correction of the image by applying an affine transform of each diffusion volume to the b0 volume and rotating b-vectors using FSL toolbox (FSL, http://www.fmrib.ox.ac.uk/fsl/). After the diffusion tensor and its eigenvector was estimated for every voxel, we applied a deterministic tractography algorithm (35) initiating a single streamline from the centre of each voxel. Tracking was stopped when the angle change was too large (35 degree of angle threshold) or when tracking reached a voxel with a fractional anisotropy value of less than 0.2 (37).

The centre coordinates of each voxel were the start of a single streamline, the total number of streamlines never exceeded the number of seed voxels. The number of connecting streamlines were used to determine the connectivity matrix (S), as the streamline count has recently been confirmed to provide a realistic estimate of white matter pathway projection strength (38). Distance matrices were also constructed using the mean fibre length of the streamlines connecting each pair of ROIs (Figure 4). The surface area of each ROI was found using FreeSurfer for cortical regions and for subcortical areas by computing the interface area to the white matter in T1 space (39).

**Figure 4.**
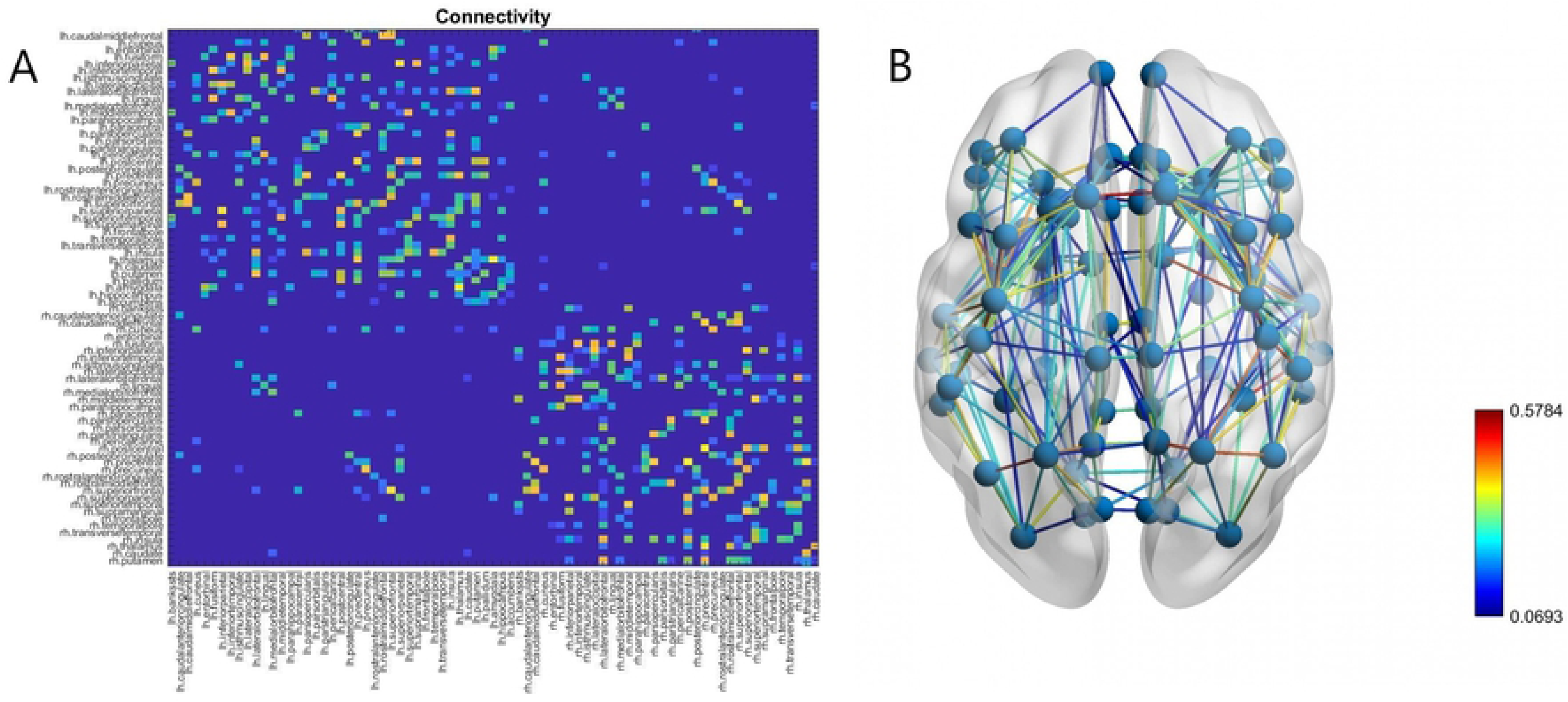
The connectivity matrix (A) obtained by the process described in the section Patient data for a healthy subject and (B) the network of nodes representing the brain of that subject. The weight of each connection (derived from the number of streamline counts between regions) is indicated by its colour.

### Modified Wilson-Cowan Model

Our model consists of a network of 82 coupled modified Wilson-Cowan oscillators, each representing a single brain region. In order to include divisive inhibition into our model, each W-C node consists of one excitatory and two inhibitory populations (Figure 5). The first inhibitory population represents interneurons firing at the dendrites of the postsynaptic neurons (subtractive inhibition) and the second inhibitory population represents interneurons firing directly at the soma of the postsynaptic neurons, delivering divisive inhibition. For the implementation of the model we followed the methodology and notation of (40). All the notations that we use for the description of the model is summarised in table 1.

**Figure 5.**
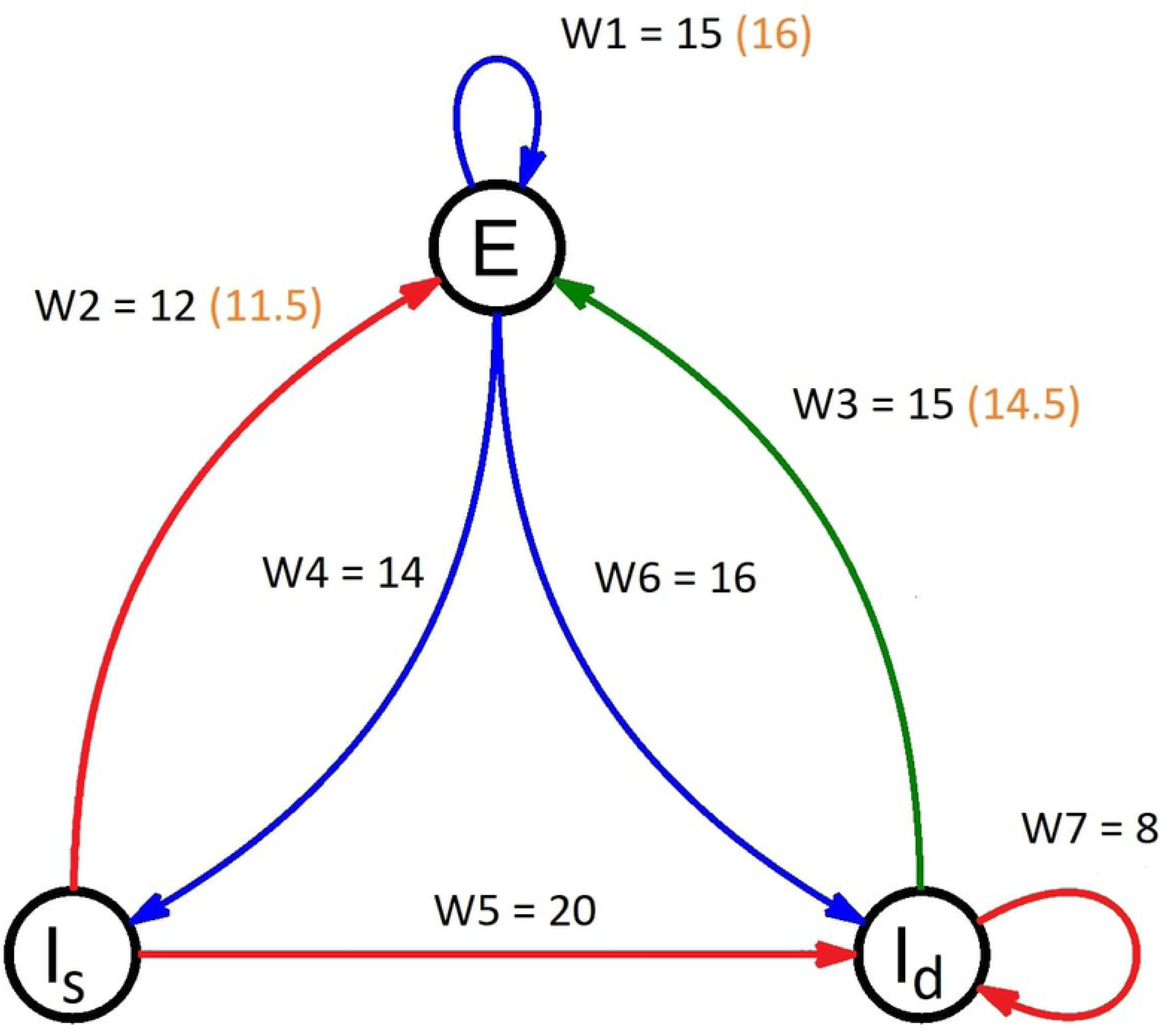
A diagram of a Wilson-Cowan node used in the model. The blue arrows indicate an excitatory connection while the red and green arrows indicate subtractive and divisive inhibitory connections respectively. The weights of each connection are indicated above every arrow. The numbers in the orange parentheses are the weight values that differ for the stimulated (epileptogenic) regions in the epileptic patients.

Of course, the model described in (40) has been designed to simulate the connectivity of a cortical microcircuit and not the connectivity of sub-cortical regions. Still, a number of studies (41-43) have shown the presence of shunting inhibition (in addition to regular subtractive inhibition) in many of the subcortical areas we used in our study. Thus, we felt that the inclusion of both inhibitory populations in the nodes representing subcortical regions was justified.

According to this model, the activity of each region is modelled by the following delayed differential equations (DDE’S):

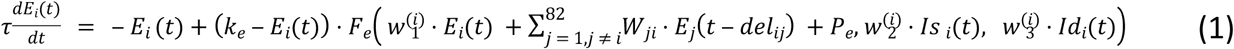

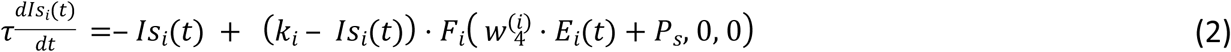

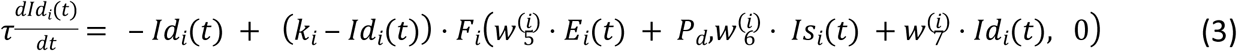

In order to account for the divisive inhibition a modified input-output function is required:

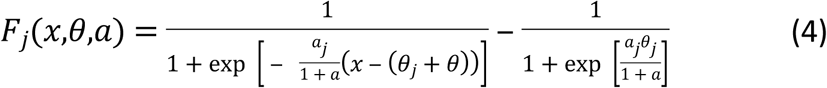

For, *j* ∈ {*e, i*} where *e* stands for excitatory and *i* stands for inhibitory. The inhibitory populations have the same input-output function and the same constants since they are assumed to respond to inputs in a similar way. However, the difference in the type of inhibition those neurons deliver to the excitatory population is due to their different targeting onto the postsynaptic neurons, that is, somatic vs dendritic.

The constant *k*_*j*_, *j* ∈ {*e,i*} is given by:

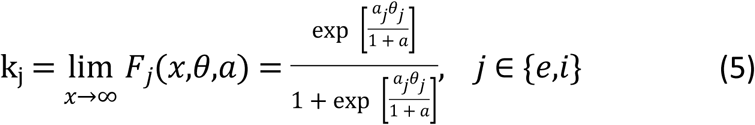

As is the case with the sigmoid function the constant *k*_*j*_ is the same for both inhibitory populations In our study, the constants of the sigmoid were set at *θ*_*e*_ = 4, *θ*_*i*_ = 3.7, *a*_*e*_ = 1.3, *a*_*i*_ = 2, following the values used at (40). Moreover, the external inputs of the inhibitory populations were set to *P*_*s*_ = *P*_*d*_ = 1 while the input of the excitatory population was set to *P*_*e*_ = 2. Other values were considered for *P*_*e*_ ranging from 1.1 to 4 (the range where the system produces oscillations) with results similar to the ones presented here. Providing no input to the inhibitory populations (*P*_*s*_ = *P*_*d*_ = 0) results in a lack of long term stable oscillations and therefore we restricted the parameter value to *P* > 0. A detailed description of all notation used is given in table 2.

**Table 2.** Notation used in the text and interpretation.

| Notation | Interpretation |
| --- | --- |
| $E_i(t)$ | Activity of the excitatory population of region i at time t |
| $Is_i(t)$ | Activity of the subtractive inhibitory population of region i at time t |
| $Id_i(t)$ | Activity of the divisive inhibitory population of region i at time t |
| $w_k^{(i)}$ | Weight of the k-th connection of region i |
| $W_{ij}$ | Weight of the connection between regions i and j |
| $del_{ij}$ | Time delay between regions i and j |
| $P_e$ | External input of the excitatory population |
| $P_s$ | External input of the subtractive inhibitory population |
| $P_d$ | External input of the divisive inhibitory population |
| $F_e(x, \theta, a)$ | Sigmoid function for the excitatory population |
| $F_i(x, \theta, a)$ | Sigmoid function for the Inhibitory populations |
| $\theta$ | Variable of the sigmoid representing subtractive modulation |
| $a$ | Variable of the sigmoid representing divisive modulation |
| $\theta_e$ | Minimum displacement in case no subtractive inhibition is delivered to the excitatory population |
| $a_e$ | Maximum slope in case no divisive inhibition is delivered to the excitatory population |
| $\theta_i$ | Minimum displacement in case no subtractive inhibition is delivered to the inhibitory populations |
| $a_i$ | Maximum slope in case no divisive inhibition is delivered to the inhibitory populations |
| $k_e$ | Constant for the excitatory population |
| $k_i$ | Constant for the inhibitory populations |

### Connectivity and Plasticity

The weights *W*_*ij*_ between brain regions were initialized according to the brain anatomy of each patient using the data described in the section ‘‘Patient data’’. Specifically, given the matrix *S* of the streamline counts for an individual subject we followed the original study (31) and initialised the connectivity matrix *M* as:

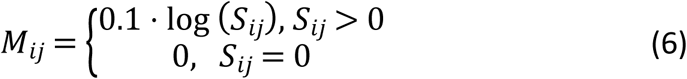

During the simulation, the weights were updated every 10 milliseconds by the following learning rule:

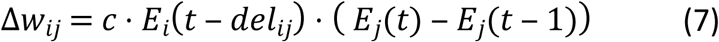

We chose this simple rule in order to represent the effects of spike timing dependent plasticity (44) in neuron populations. The learning rate was set at c = 0.1. Other values were considered, and similar results were obtained with the only difference being the speed of weight change. Still, the pattern of activity remained the same for all the values we examined as can be seen in figure 6.

**Figure 6.**
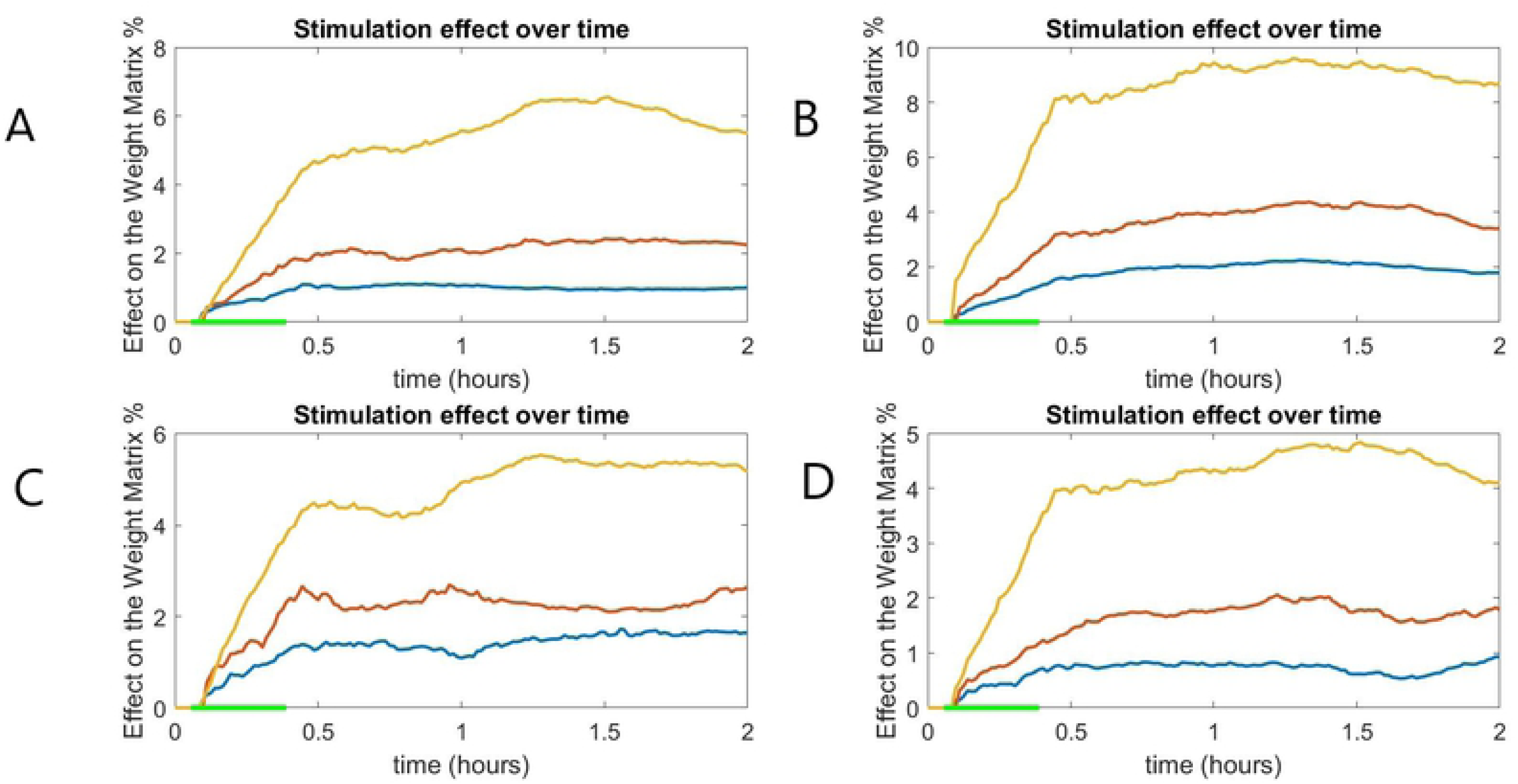
The global connectivity difference measures two epileptic (A,B) and two healthy (C,D) subjects for different learning rates: c = 0.05 (blue), c = 0.1 (red) and c = 0.2 (yellow). The effect of stimulation on the global connectivity is different depending on the learning rate but the overall pattern remains similar. The green line at the x-axis indicates the period of stimulation.

The weight matrix was normalised after each update (45, 46) by the following rule:

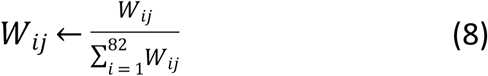

For the internal weights 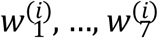 of each region we used two different sets of initial values. The first set of values was chosen to represent the connectivity of a healthy brain region while the second set was chosen to represent an epileptogenic region. The values of the healthy region were decided after an extensive parameter search, starting at the values used by (40) and examining values between 8 and 21 (the range at which the system produces oscillations). The values we selected lead to high amplitude oscillations in all three populations during the first hours of the simulation. The amplitude of the oscillations gradually decreases and stabilizes after some hours. It must be noted that the final values were chosen to facilitate the dynamics of the system and may not correspond to the connectivity of a real biological system. Still, using different parameters usually resulted in oscillations of different amplitude and consequently slower stabilization periods, but as a general rule did not lead to radically different behaviour in the system.

After the values of the healthy region were established, the values of the epileptogenic regions were derived by increasing the weights of excitatory connections and reducing the weights of the inhibitory connections. Those changes aimed at increasing the excitability of those regions (increased excitatory and decreased inhibitory input) in order to simulate the dynamics associated with epilepsy. The difference in behaviour of the epileptogenic regions was small but observable (oscillations of increased amplitude and occasional seizure-like activity when the input to their excitatory regions was increased), as with the original connection eights, choosing different values led to slightly different results (the more excitable the regions, the greater the effect of stimulation), but the main observations remained the same. The values chosen are presented in Figure 5

The weights *w*_1_, *w*_2_, *w*_3_, *w*_4_, *w*_6_ were updated every 10 milliseconds according to a modified version of the rule we used for the external connections with subsequent normalization after every update.

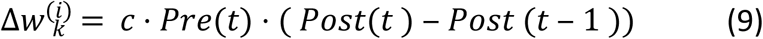

Where *Pre*(*t*), *Post*(*t*) are the activities of the presynaptic and the postsynaptic populations, respectively. Several proposed mechanism of internal plasticity were considered, but due to the lack of a consensus about a general mechanism of inhibitory plasticity (44, 47)—especially in neural mass models—we chose to use this simple intuitive rule, similar to the rule we used for the external connections. The most commonly used learning rule for inhibitory plasticity, introduced in (48) could not be used in this model due to long term instability in the networks dynamics.

For the normalization, we employed the same rule used for the global connectivity:

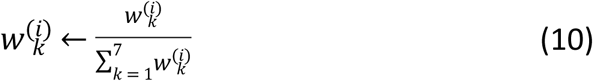

Since there has been little research on how inhibitory to inhibitory plasticity could be implemented in a neural mass model, the weights *w*_5_ and *w*_7_ were kept stable. The learning rate was set at c = 0.05.

Finally, the delays were initialized for each patient, as the length of the fibres connecting two regions divided by the speed of spike propagation. For the calculation of the delays we considered all axons to be myelinated and thus the spike propagation speed was set at 7 m/s (49, 50). To calculate the distance between regions, we selected the fibre trajectory length—which we calculated using deterministic tracking of diffusion tensor imaging data—instead of the Euclidian distance in order for the delays to be more biologically realistic.

### Stimulation

Each session of stimulation was modelled as a decrease of 50% (the stimulation is cathodal, due to better reported experimental results (10)) in the external input of the three regions (amygdala, hippocampus and parahippocampal gyrus) most commonly responsible for seizures in these patients, for a period of 30 minutes. Despite two of these regions being sub-cortical, the ability of transcranial stimulation to affect them has been demonstrated in past studies (51-53). Stimulation in all cases started at t =200s after the beginning of the simulation. This initial period was allowed for the oscillations of the system to stabilize before stimulation begins.

The choice of stimulation parameters was made in order for the model to correspond to a working protocol of TCS (54). Due to the computational constraints of such large simulations (55), we modelled only one session and an additional resting period of 24 hours.

### Model Implementation

The model was initialized with the data of each patient as described in the previous sections and two simulations–with and without stimulation-run in parallel for a period of 24 hours with snapshots of the weight matrices taken every 50 seconds. The large system of DDE’s (246 equations) was solved by using Matlab’s dde23 delayed differential equation solver.

The effect of the stimulation on the connectivity at every time step was measured in the following ways:

1. The global effect of the stimulation on the connectivity of the brain was measured as the difference (%) of the connectivity matrices *M* = (*W*_*ij*_):

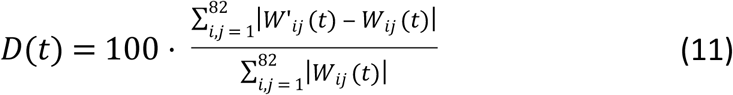

where *W*’_*ij*_ is the weight between regions *i* and *j* at time t after stimulation and *W*_*ij*_ (*t*) is the weight between regions *i* and *j* at time t without stimulation. This measure represents the effect stimulation has on the inter-region connections of the brain.

2. The effect of the stimulation on the internal connectivity of each region (local effect) was measured as the difference (%) of the internal weights in the stimulated and the non-stimulated versions:

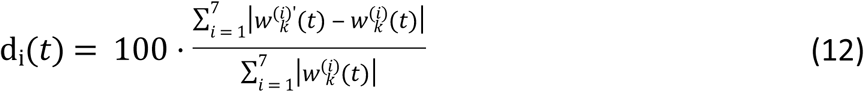

where i =1,…,82 the brain region 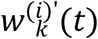 is the k-th weight of the i-th region at time t in the stimulated version and 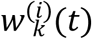 is the i-th weight of the k-th region at time t in the non-stimulated version. These measures represent the effect of stimulation on the internal connectivity of each region.

### Connectivity measure

In order to study the effect of stimulation on the regions that received no direct stimulation, we examined several connectivity metrics that could explain such an effect. One of those metrics is the Jaccard index. The Jaccard index of two regions measures the similarity in connectivity (the common neighbours) and is defined as:

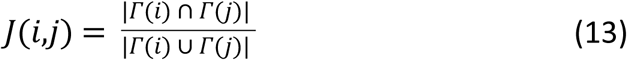

Where *Γ*(*i*) is the set of nodes connected to node *i* and |*A*| is the number of elements of the set *A*. In our study, we defined the Jaccard index of a secondary region *i* to be:

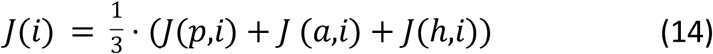

Where *p,a,h* are the stimulated regions.

### Clinical Significance

In the end of our study we present a hypothetical test that aims to the outcome of surgery. In order to assess the effectiveness of a clinical test the following measures are used (56):

1. 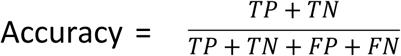, which measures the ability of the test to differentiate the patient and healthy cases correctly.
2. 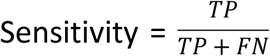, which measure the ability of the test to determine the patient cases correctly
3. 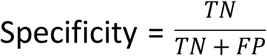, which measures the ability of the test to determine the healthy cases correctly

Where TP means true positive, TN means true negative, FP means false positive and FN means false negative.

